# Tree diversity and the temporal stability of mountain forest productivity: testing the effect of species composition, through asynchrony and overyielding

**DOI:** 10.1101/2020.01.20.912964

**Authors:** Marion Jourdan, Christian Piedallu, Jonas Baudry, Xavier Morin

## Abstract

Climate change modifies ecosystem processes directly through its effect on environmental conditions, but also indirectly by changing community composition. Theoretical studies and grassland experiments suggest that diversity may increase and stabilize communities’ productivity over time. Few recent studies on forest ecosystems suggested the same pattern but with a larger variability between the results. In this paper, we aimed to test stabilizing diversity effect for two kinds of mixtures (*Fagus sylvatica* - *Quercus pubescens* and *Fagus sylvatica* - *Abies alba*), and to assess how climate may affect the patterns. We used tree ring data from forest plots distributed along a latitudinal gradient across French Alps. We found that diversity effect on stability in productivity varies with stand composition. Most beech–fir stands showed a greater stability in productivity over time than monocultures, while beech–oak stands showed a less stable productivity. Considering non-additive effects, no significant trends were found, regardless the type of mixed stands considered. We further highlighted that these patterns could be partially explained by asynchrony between species responses to annual climatic conditions (notably to variation in temperature or precipitation), overyielding, and climatic conditions. We also showed that the intensity of the diversity effect on stability varies along the ecological gradient, consistently with the stress gradient hypothesis for beech-oak forests, but not for beech-fir forests. This study showed the importance of the species identity on the relationships between diversity, climate and stability of forest productivity. Better depicting diversity and composition effects on forest ecosystem functioning appears to be crucial for forest managers to promote forest adaptation and maintain timber resource in the context of on-going climate change.

## 1 INTRODUCTION

Climate change affects ecosystem functioning and its related services directly through its effect on environmental conditions (Olesen et al. 2007; Malhi et al. 2008), but also indirectly by changing community composition (Bertrand et al. 2011), because changes in community diversity may affect ecosystem functioning (Hooper et al. 2005; Cardinale et al. 2012), as shown in many ecosystems for mean productivity (Hector 1999; Paquette and Messier 2011). Predicting the direction and strength of these two types of effects (direct and indirect) remains a difficult task (Morin et al. 2018), partly because indirect effects have multiple facets. Former studies have shown that species diversity affects ecosystem productivity through a “performance enhancing” effect (i.e. an increasing mean ecosystem productivity) and a “buffering effect” (i.e. a decrease in temporal variance of ecosystem productivity). The latter effect is consistent with the ecological insurance hypothesis (Yachi and Loreau 1999) stating that diversity should mitigate the impact of environmental fluctuations on ecosystem functioning. Thus, changes in species composition within communities seems to not only affect mean productivity, but also temporal stability - often assessed as the inverse of the coefficient of variation of productivity over time (μ/σ, with µ and σ being respectively the mean and standard deviation of the time-series of the considered process Lehman and Tilman (2000)). Regarding the performance enhancing effect, numerous theoretical, empirical and experimental studies have shown that species-rich communities generally benefit from “overyielding” in comparison to species-poor communities (Cardinale et al. 2007; Loreau and Hector 2001). It means that productivity is greater-than-expected in diverse ecosystems when compared to expectations inferred from component monocultures. This pattern has been mostly explained by “niche partitioning” in more diverse communities, i.e. a greater complementarity between species niches leads to an increase in resource uptake at the community level and also by decreased competition in more diverse communities, especially between conspecifics (Tilman 1999; Hooper et al. 2005; Jucker et al. 2014). Experimental results on this issue mainly come from herbaceous communities, but overyielding patterns was also shown in tree communities (Jones, McNamara, & Mason, 2005; Pretzsch, 2005) provided by empirical observations (Paquette and Messier 2011; Vilà et al. 2013; Toïgo et al. 2015). These studies show, on average, a positive effect of diversity on forest productivity which may strongly depend on site fertility (Pretzsch et al. 2015), with notably stronger effects in harsher environment in comparison with lowlands (Paquette and Messier 2011; Toïgo et al. 2015; Jactel et al. 2018) or on species composition (Forrester 2014).

The buffering effect is much less explored and tested, particularly in forests. Here, we focus on temporal stability (*TS*) of ecosystem productivity, calculated using annual basal area increment (*BAI*), across years (Tilman 1996). A positive effect of biodiversity on *TS* is theoretically expected, as shown by analytical or simulation models for various ecosystems (Morin et al. 2011; Loreau and de Mazancourt 2013). This expectation for diversity-stability relationships is confirmed through experiments (Hector et al. 2010; Isbell et al. 2015) and observation-based studies (Isbell et al. 2009; Hautier et al. 2014), mostly in grassland ecosystems. Only few studies deal with *TS* of stand productivity in forest ecosystems (DeClerck et al. 2006; Jucker et al. 2014; Aussenac et al. 2017; del Río et al. 2017). Most studies based on observation *in situ* show a positive effect of diversity on *TS* in forest ecosystems (DeClerck et al. 2006; del Río et al. 2017), except Jucker et al. (2014) who find positive or negative effects depending on study sites (while the effect of site characteristics, e.g. soil fertility, climate conditions, were not tested). To conclude there is not yet strong consensus on tree species richness effect on *TS* of forest stands. More studies are necessary to properly depict diversity effects on temporal stability of productivity.

Theoretically diversity effect on temporal stability of ecosystem processes remain weakly understood (Loreau and de Mazancourt 2013). Two main mechanisms are proposed: i) overyielding and ii) asynchrony in species responses to environmental conditions (Hector et al. 2010). Overyielding mathematically increase ecosystem *TS* by increasing mean ecosystem productivity (higher *µ*) while keeping the temporal variability of ecosystem processes (*σ*) constant (Lehman and Tilman 2000). The effect of asynchrony is related to the temporal complementarity between species, occurring in the case of differential responses of species to environmental conditions, and reducing variability in productivity at the community level (Loreau and de Mazancourt 2013) – induced by niche differences among species (Yachi and Loreau 1999). In forest ecosystems, the few studies that have focused on this issue found that stability in productivity generally increased with asynchrony, either through modeling (Morin et al. 2014) or empirical approaches (Jucker et al. 2014; del Río et al. 2017).

Furthermore, the impact of climate conditions on the strength of the diversity-stability in productivity relationships remains largely unknown, especially in tree communities. To our knowledge, only Jucker et al. (2014) or del Río et al. (2017) explored this effect and found that environmental conditions may affect the stability of aboveground wood production. However, this finding is not obtained by comparing the same species mixtures along climatic gradients (Jucker et al. 2014) or is not explicitly tested (del Río et al. 2017). Yet, depicting how climate conditions may impact diversity effect on forest functioning appears critical in the context of on-going climate change, especially in Mediterranean and/or mountainous environments which are particularly sensitive to future environmental changes (Thuiller et al. 2005). Differences between the responses of species are supposed to be more marked in stressful conditions, according the stress gradient hypothesis (*SGH*, Lortie and Callaway 2006; Maestre et al. 2009), thus exacerbating the diversity effect in such conditions (Morin et al. 2018).

Here we aim at testing whether wood productivity (measured using *BAI*) at the stand level is more stable over time in mixed stands than in monospecific stands and whether climate conditions may affect this possible diversity effect. To do so, we focus on two types of mixed-species forest, beech (*Fagus sylvatica* [L.]) – pubescent oak (*Quercus pubescens* [L.]) and beech – silver fir (*Abies alba* [L.]) forests in the French Alps. These mountain forest types are distributed along strong latitudinal climatic gradients (temperature and precipitation gradient), which makes them particularly interesting to explore climate effects on ecosystem functioning. Furthermore, these tree species and forest types are particularly important in this region, for both ecological and economic reasons. For instance, beech-oak forests are considered of strong ecological interest in Southern French Alps (Regnery et al. 2013), while silver fir is a central species of the Northern Alps timber industry. They are thus particularly interesting to study because : *i)* these forests have been identified as especially vulnerable to climate change (Courbaud et al. 2011), *ii)* beech-fir forests are well represented ecosystems in Northern French Alps while beech-oak forests are present in the Southern part, and *iii)* it allowed to compare two mixtures with contrasting composition. More precisely, these mixtures include two broadleaf species with contrasting shade tolerance (beech being more tolerant than pubescent oak) and drought tolerance (beech being less tolerant than pubescent oak) and a conifer and a broadleaf species with similar shade tolerance and contrasted drought tolerance (beech being more tolerant than silver fir, Niinemets and Valladares 2006). These contrasting differences in functional properties of the two kinds of mixtures may induce various responses to same climate conditions between species. For instance, we expect that more shade-tolerant species may benefit from mixing with a less shade-tolerant species (Toïgo et al. 2015, 2017), i.e. more shade-tolerant species may experience weaker competition in this case.

In this study, we test whether the productivity of mixed stands is more stable than the productivity of monospecific ones by using a network of field plots organized by triplets (i.e. a 2-species mixed stand and the two respective monospecific stands). We specifically aim at answering to the following questions:

i. Is the temporal stability of productivity at the plot scale greater in mixed stands than in monospecific stands? We expect that temporal stability of forest productivity should be stronger in mixed stands than in monospecific stands, with a stronger effect under more stressful conditions.
ii. Does species identity and climate conditions affect the direction and magnitude of the diversity effect on temporal stability in productivity? We expect the magnitude of the diversity effect to be dependent on species identity and that the role of asynchrony should increase with increasing environmental harshness, according to the SGH.
iii. What is the relative importance of the roles of overyielding *vs.* asynchrony on temporal stability? We expect that asynchrony should affect stability in productivity more strongly than overyielding, according to theoretical (Morin et al. 2014) and empirical findings (Cardinale et al. 2013).

## 2 MATERIALS AND METHODS

### 2.1 Field design

The field design is distributed along a six-sites latitudinal gradient in the French Alps with contrasting climatic conditions - from North to South: Bauges, Vercors, Mont Ventoux, Luberon-Lagarde, Grand Luberon and Sainte-Baume (Fig. 1 and Table S1, see also Jourdan et al. 2019). Sites were selected to minimize topographical conditions variability. On each site, triplets of plots were distributed along elevational gradients. All sites were characterized by limestone bedrock, with a North to West aspect.

**Figure 1.**
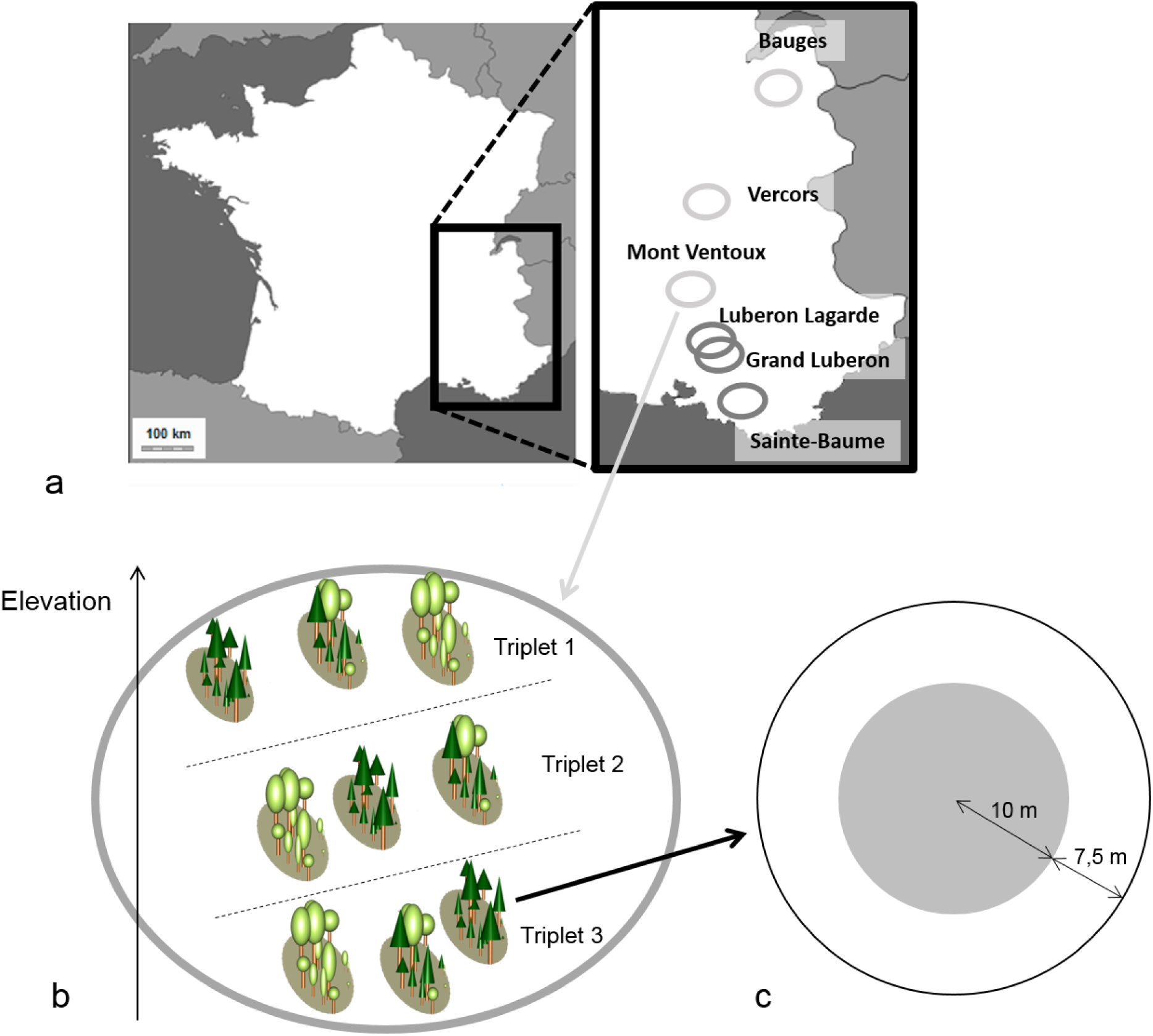
Field design description. (a) Study area and sites where plots were sampled. Beech-fir stands (light grey circles) are Mont Ventoux, Vercors and Bauges, with plots sampled in beech-fir forests. Beech-oak forests (dark gray circles) are in Luberon Lagarde, Grand Luberon and Sainte-Baume, with plots sampled in beech-oak forests. (b) Schematic representation of a site with three triplets stands (one monospecific stand of species A (beech), one monospecific stand of species B (fir or oak), and one mixed stand with species A and B) distributed along an elevational gradient. (c) Schematic representation of a plot, with an inner circle (grey area) in which all trees with a DBH > 7.5 cm were sampled (and an external 7.5m-crown -*buffer zone*-in which only the dominant trees). (Paint 6.1)

**Table 1:**
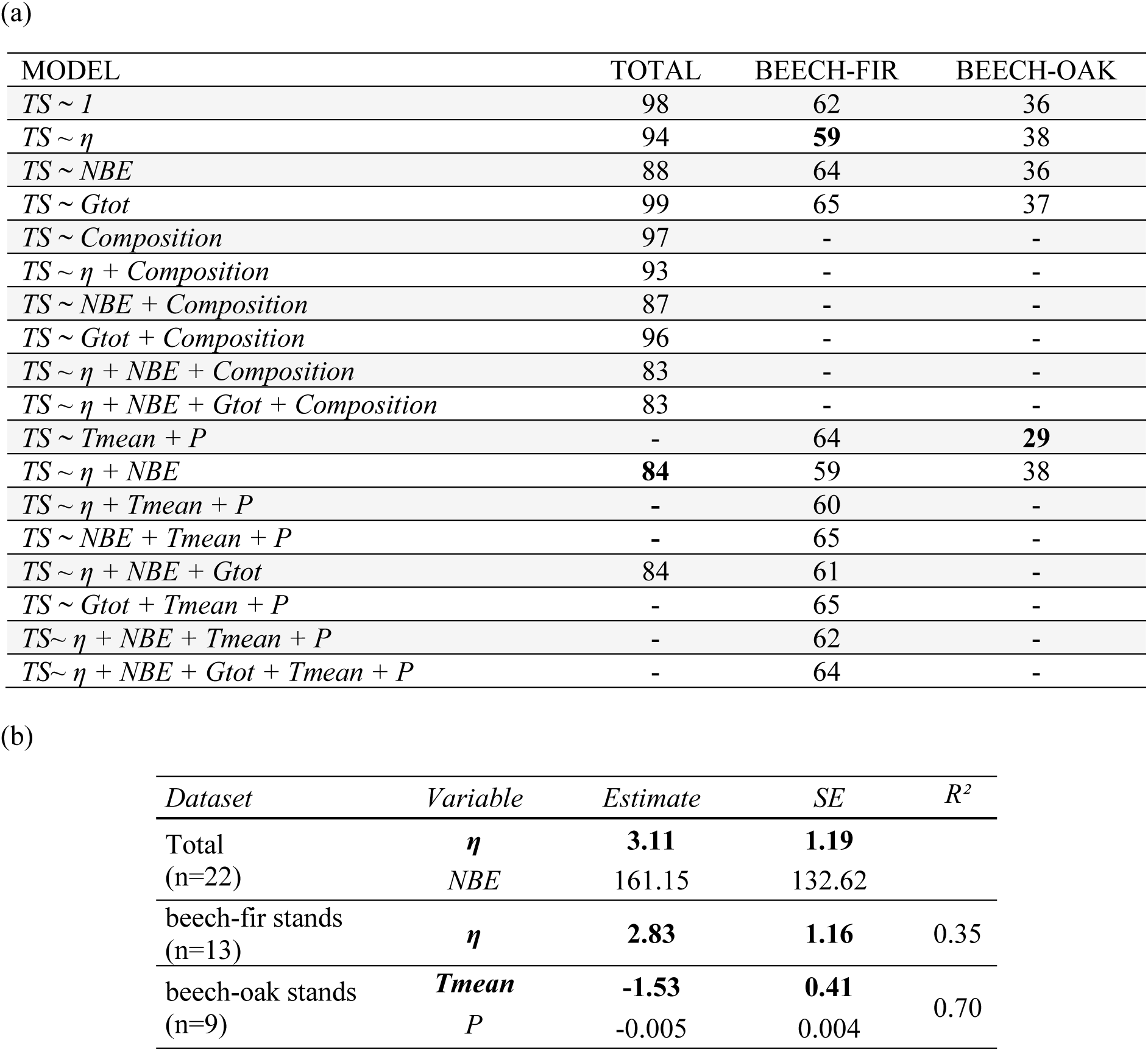
Linear mixed models of Stand Temporal Stability Linear mixed models of Stand Temporal Stability (*TS*) against asynchrony (*ƞ*), Net Biodiversity Effect (*NBE)* and climate variables for the whole dataset (*Total*), and composition are “beech-fir” in North and “beech-oak” in South. For beech-oak and beech-fir stands of the gradient taken separately, *T_mean_* and *P* are annual mean temperature average and annual precipitation average across 1997-2012 respectively. In every model *site* is a random effect. *n* represents number of triplets considered in each model. (a) AIC the most relevant tested linear mixed models. AIC of most parsimonious models are highlighted in bold. (b) Coefficients estimates of explanatory variables in the selected models. Significant explicative variables are in bold.

Northern sites (Bauges, Vercors, Mont Ventoux) are composed of beech-fir forests and Southern sites (Luberon Lagarde, Grand Luberon, Sainte-Baume) are composed of beech-oak forests. Stand structure was high forest, except in Grand Luberon where all stands are coppices. The plots were grouped into dense triplets within a site, i.e. the combination of a beech monospecific plot, a fir or oak monospecific plot and a mixed plot (fir-beech or oak-beech). These triplets were distributed along an elevational gradient at the site level. Focusing on forest mixed plots with two species allows testing complementary effects in a more precise way (Forrester and Bauhus 2016; Aussenac et al. 2017) and better identifying explanatory mechanisms at a local scale. A total of 67 plots were sampled across the design (Table S1), organized in 22 triplets (one plot was not used in this study).

A plot was composed of a 10 m-radius disk of homogenous stand structure and composition (Fig. 1). We measured tree characteristics: species identity, localization, height, crown depth and diameter at breast height [DBH], i.e. at 1,30 m. We sampled one core per tree at breast height for dendrochronological analyses using a Pressler borer, i.e. all trees with a DBH larger than 7.5 cm were cored, except for coppice stands in which only the largest stem of each coppice was cored. We thus sampled all the trees regardless of their status (dominant or understory). The slope, elevation and aspect were measured for each plot.

### 2.2 Climate data

To quantify the effect of climate on tree growth, we first selected variables often used in dendrochronological studies according to their known effect on tree growth (Cailleret and Davi 2010): mean annual temperature and sum of annual precipitation, both averaged across the study period (1997 to 2012). We chose only two climatic variables to allow simple result interpretation and to keep enough statistical power for linear models. Monthly values of precipitation and temperatures were extracted from 1 km-resolution GIS layers at the whole national level (Piedallu et al. 2016). These maps were created using data from 119 and 214 non-interrupted weather stations from the Météo France network, for precipitation and temperature respectively. To build the climatic maps, a monthly model was created for each variable using Geographically Weighted Regression (GWR, Fotheringham et al., 2002) with spatially distributed variables describing topography, solar radiation, land use and distances to the seas (Piedallu et al, 2016). Cross-validation was used to validate these maps, showing on French territory an average r² ranging from 0.80 for precipitation to 0.94 for mean temperatures.

### 2.3 Basal area increments dataset and analyses

We analyzed growth dynamics using tree rings collected over the 15 years from 1997 to 2012. Each tree ring was first photographed with a large-resolution camera coupled with a binocular lens. The width of each ring was then assessed with ImageJ software (https://imagej.nih.gov/ij/index.html), with an accuracy of 0.01 mm. All cores were cross-dated using the method published by (Cook et al 1990, Lebourgeois and Merian 2012). Diameter increments were transformed into basal area increments (*BAI*) using measured DBH (Biondi and Qeadan 2008). We obtained reliable growth time-series for 1235 trees (596 beech, 387 fir, 240 oak and 12 for other trees species – mostly Scots pine, maple and spruce trees).

This study aimed at assessing forest productivity by sampling almost all trees in a plot instead of sampling only dominant trees (Lebourgeois et al 2010, Lebourgeois et al 2013). Considering all trees should allow to better quantify the whole competitive environment within each plot. However, we did not obtain growth data for all trees due to the difficulty of reading some cores, especially in the southern sites, because of very narrow rings or difficulty to distinguish rings. As the unreadable cores were not equally distributed across the network of plots but also inside each triplet, it was not possible to focus on only the subsample of available trees (i.e. trees with a usable core) to calculate *TS* for the whole plot. Hence to assess the productivity of the whole plot, we reconstructed of the temporal series of *BAI* of the missing individuals, i.e. 924 trees – thus *c.a.* 40% of total dataset. To do so, we fitted a linear model for each stand type and each species in each site for each year, with *BAI* as the variable to explain and annual tree basal area as explanatory variable. With the model’s estimates, we predicted a *BAI* for each missing tree time-series (models’ estimates shown in Table S2).

**Table 2:** Linear mixed models of Net Biodiversity Effect on Stability Linear mixed models of Net Biodiversity Effect on Stability (*SNBE*) against asynchrony (*ƞ*), Net Biodiversity Effect (*NBE)* and climate variables for the whole dataset (*Total*), and composition are “beech-fir” in North and “beech-oak” in South. For beech-oak and beech-fir stands of the gradient taken separately, *T_mean_* and *P* are annual mean temperature average and annual precipitation average across 1997-2012 respectively. In every model *site* is a random effect. *n* represents number of triplets considered in each model. (a) AIC the most relevant tested linear mixed models. AIC of most parsimonious models are highlighted in bold. (b) Coefficients estimates of explanatory variables in the selected models. Significant explicative variables are in bold.

| MODEL | TOTAL | BEECH-FIR | BEECH-OAK |
| --- | --- | --- | --- |
| *SNBE ~ 1* | 107 | **68** | **21** |
| *SNBE ~ $\eta$* | 103 | 68 | 21 |
| *SNBE ~ NBE* | 97 | 70 | 22 |
| *SNBE ~ $\eta$ + NBE* | **92** | 70 | 23 |
| *SNBE ~ Composition* | 105 | - | - |
| *SNBE ~ $\eta$ + Composition* | 101 | - | - |
| *SNBE ~ NBE + Composition* | 94 | - | - |
| *SNBE ~ Gtot + Composition* | 104 | - | - |
| *SNBE ~ $\eta$ + NBE + Composition* | 91 | - | - |
| *SNBE ~ $\eta$ + NBE + Gtot + Composition* | 90 | - | - |
| *SNBE ~ Tmean + P* | - | 71 | 22 |
| *SNBE ~ $\eta$ + NBE + Tmean + P* | - | 72 | - |

(b)
| Dataset | Variables | Estimate | SE | $R^2$ |
| --- | --- | --- | --- | --- |
| Total<br>(n=22) | **$\eta$** | **3.03** | **1.47** |  |
|  | *NBE* | 126 | 163 |  |
| beech-fir stands<br>(n=13) | - | - | - | - |
| beech-oak stands<br>(n=9) | - | - | - | - |

In this study we used annual *BAI* at the plot level over the last 15 years. The increment at year *j* (*BAI_j_*) was calculated by summing the *BAI* of all trees for each plot:

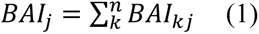

with *n* being the number of trees in the plot and *BAI_kj_* being the basal area increment of tree *k* at year *j*.

### 2.4 Temporal stability: Climate conditions, asynchrony and overyielding effects

We defined the inverse of coefficient of variation of basal area increment (Lehman and Tilman 2000) to calculate *TS* from the growth time-series:

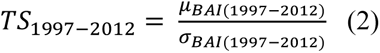

with *μ_BAI_* and *σ_BAI_* being respectively the mean and standard deviation of plot annual basal area increments between 1997 and 2012. A high *TS* value corresponds to a large temporal stability in productivity.

Two main diversity-mechanisms are supposed to affect temporal stability of plot productivity: change in mean productivity across years (hereafter called “overyielding” effect) and asynchrony in species response to environmental conditions (hereafter called “asynchrony”). To detect an overyielding effect, we calculated the net diversity effect *NBE_i_*, (Loreau and Hector 2001) for each mixed stand *i*, as follows:

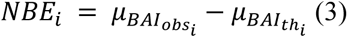

with *μ_BAI_obs_i___* and *μ_BAI_th_i___* being the observed and theoretical averaged annual productivity respectively between 1997 and 2012. A positive *NBE* means that mixed stand productivity was larger than expected when interspecific interactions are discarded (Loreau and Hector 2001).

For each mixed stand, we calculated an index of asynchrony to test for its possible stabilizing effect on *BAI* (de Mazancourt et al. 2013). We used the metric *η* developed by Gross et al. (Gross et al. 2014), defined as the average across species of correlations between *BAI* time-series of each species and *BAI* time-series of all other species in the community. Therefore, *η* is calculated as follows:

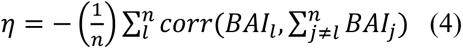

with *BAI_l_* being the productivity of species *l* in a community of *n* species. The index ɳ ranges from −1 when species are perfectly synchronized to 1 when species are perfectly asynchronized, as defined in (Gross et al. 2014). The case ɳ = 0 occurs when species’ growths fluctuate independently. In our study, the maximum diversity is *n =* 2, thus ɳ can be simply written as:

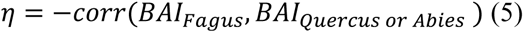

with *BAI_Fagus or Quercus or Abies_* being the productivity of each species in the same triplet.

We first tested whether *TS* of mixed stands varied across stand composition, for the different parts of the gradient. *TS_i_* were tested using linear mixed models and AIC comparisons. We used the following model for the whole gradient, for each plot *i*:

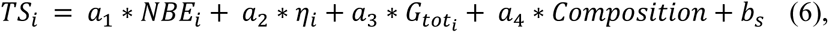

with *NBE_i_* representing net biodiversity effect in the mixed plot *i*, *η_i_* representing asynchrony, *G_tot_i__* representing total basal area of mixed plot, for each triplet, *Composition* representing the kind of mixed stand considered (i.e. fir-beech forest for North and oak-beech forest for South) and a_1->4_ are the respective fitted coefficients. *b_s_* is site effect (defined as a random effect).

As the effects of climate and stand composition may be confounded along the whole gradient, we thus used two other models to consider the climatic conditions of site effect on mixed plot i (instead of a random effect for the site), one for each sub-gradient (i.e. beech-fir and beech-oak forests separately):

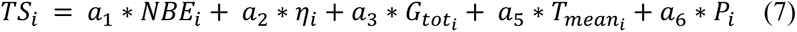

with *T_mean_i__* being the mean annual temperature and *P_i_* being the sum of annual precipitation and a_5_ and a_6_ are the respective fitted coefficients.

To test whether *TS* between mixed and monospecific stands may change with environmental conditions, i.e. testing the SGH with our mixed-pure stand comparison, we focused on mixed stands *vs.* beech monospecific stands because beech was the only species present along the whole gradient. To do so, we calculated the ratio between temporal stability of mixed and monospecific forests 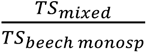 This ratio was superior to 1 in 13 cases among the 22 triplets. Then we carried-out a linear mixed model of the ratio against the mean productivity (in m².ha^-1^.yr^-1^) of each mixed stand, i.e. using the mean productivity as an integrative proxy for site’s conditions.

All analyses have been carried-out with R software (R version 3.3.0). Each model residual distribution follows the assumptions of normality and homoscedasticity.

### 2.5 Non-additive effect of diversity on temporal stability

Then we tested the non-additive effect of diversity (through the net biodiversity effect, NBE) on TS, i.e. the part of mixture effect not related to mass effect of species addition on total productivity average. We quantified NBE on temporal stability of productivity (hereafter “*SNBE*”, Jourdan et al. 2019), built in analogy to NBE usually calculated for mean productivity (Loreau and Hector 2001). The *SNBE* (Eq 8) thus relied on the comparison of the observed *TS* of a mixed stand productivity *i* (*TS_obs_i__*) with a corresponding theoretical *TS* of mixed stand productivity (*TS_th_i__*). The theoretical *TS* of each mixed plot *i* was estimated for each species using the annual productivity of monospecific stands in the same triplet.

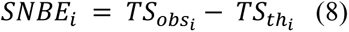

This method allowed to calculate increases or decreases in stability of productivity between mixed and monospecific stands, considering the various relative abundances of species across sites. A *SNBE* value close to 0 means that the stability in productivity of mixed stands is close to the stability expected from monospecific stands under the null hypothesis that there is no effect of interspecific interactions between co-existing species on tree growth. Positive and negative *SNBE* values respectively correspond to a larger and weaker stability of mixed stands productivity than expected from monospecific stands due to non-additive effect. We used two different approaches to compute *TS_th_i__*.

In the first approach, the expected stability of mixed stand productivity was reconstituted using species productivity in monospecific stands and the species’ relative abundance in the observed mixed stands (based on basal area). Thus *TS_th_i__* was computed using a theoretical productivity *BAI_th_i,j__* for each mixed plot *i* and for each year *j*, according to method developed by Loreau (1998). The productivity of each mixed stand has been partitioned into three parts (see Fig. S4): productivity of beech trees, productivity of accompanying species (fir or oak) and productivity of other species. The relative abundance of each species in the mixed stand (*p_Fagus_, p_Quercus or Abies_* and *p_other_*) was calculated using the summed initial basal area of each species (i.e. in 1997). The structure of stands (i.e. total basal area) was similar within a triplet in most cases (see Table S3). We removed triplets (n = 4) from this analysis because they showed dissimilar stand characteristics (total basal area, dominant height, density). We thus finally obtained 19 triplets to analyze. As the effects of “other species” were negligible regarding their weak values of basal area, they were considered constant between monospecific and mixed stands. The theoretical annual productivity of each mixed plot *i* was estimated for each species using the annual productivity of monospecific stands (*BAI_Fagus,Quercus or Abies_i,j__*) at year *j* in the same triplet. The theoretical annual productivity *BAI_th_i,j__* was thus:

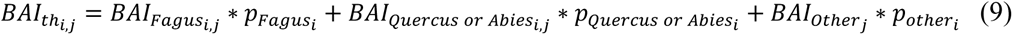

The *SNBE* of a mixed plot *i* is calculated using equations (2), (8) and (9).

Then *SNBE_i_* was calculated at the species level to better understand relative role of each species in the results of *SNBE_i_* at plot level. We compared the observed stability of productivity of only one species in each mixture with theoretical stability of productivity of same species calculated from the monospecific stand while respecting the relative abundance of species in mixture. The analysis of *SNBE_i_* at species level was done separately for each part of the gradient (i.e. beech-fir and beech-oak forests).

In a second approach, we considered the possible heterogeneity of the stands structure within triplets and thus computing *TS_th_i__* more precisely than in the first approach. To do so, we used the same methodology than in the first approach (see above) but we applied a correction factor *p_c_* for each species, to take into account the possible difference in basal area between mixed and monospecific plots of the same triplet (the results are being presented in Appendices 6 and 7):

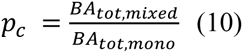

with *BA_tot,mixed_* and *BA_tot,mono_* being respectively the total initial basal area (i.e. basal area in 1997) of the mixed stand and of a monospecific stand. In this case no triplets were removed. With this approach, the theoretical annual productivity of plot *i* in the year *j* (*BAI_th_i,j__*) was thus:

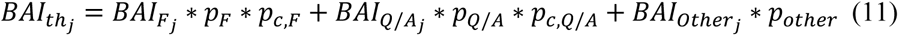

Then, for both approaches, we used a Student t-test to test whether the *SNBE* values were significantly different from 0.

We used the same kind of analyses to test for the effects of *NBE* and asynchrony on *SNBE* than for *TS*, thus using linear mixed models and AIC comparisons. We used the following model for the whole gradient (with the both approaches: with and without stand structure correction), for each mixed plot *i*:

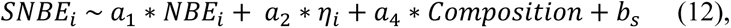

with *NBE_i_* representing net biodiversity effect, *η_i_* representing asynchrony, *Composition* representing mixture considered (i.e. “fir-beech” forest for North and “oak-beech” forest for South) and a_1,2,4_ are the respective fitted coefficients, and *b*_s_ is the site effect (defined as a random effect).

As for the tests on *TS*, we used two other models to consider the climatic conditions at the site level (instead of a random effect for the site), one for each sub-gradient:

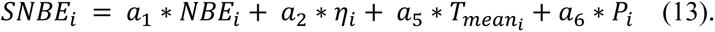

Like for the ratio of *TS* described above, we carried-out a linear model of *SNBE_i_* against the mean productivity of each mixed stand.

All analyses have been carried-out with R software (R version 3.3.0). Each model residual distribution follows the assumptions of normality and homoscedasticity.

## 3 RESULTS

### 3.1 Temporal stability (*TS*)

Temporal stability of mixed stands (beech-fir and beech-oak mixed stands combined) and monospecific stands productivity (fir, beech and oak monospecific stand combined) were not significantly different across the whole gradient (*TS*, P>0.05, t-test, n=66 plots), nor for each sub-gradients (beech-fir and beech-oak forests), although the mean *TS* was stronger for mixed stands in the beech-fir forests compare beech and fir monospecific stands (*µ_mixed_* = 7.73±0.61 (n=14) vs. *µ_monospecific_* = 6.84±0.56 (n=27), P = 0.12).

Focusing on mixed plots, we tested the effect of several potential drivers on *TS*. Relying on AIC to compare the several linear mixed models, we found that *TS* was significantly dependent on asynchrony and *NBE* for the whole gradient (Table 1-a), but only on asynchrony for the beech-fir forests. Coefficient analysis showed that *TS* increased with asynchrony between species (Table 1-b and Fig. 3-a). In beech-oak forests, coefficient of model analysis notably showed that *TS* decreased with increasing temperatures (see Table 1-b).

**Figure 2.**
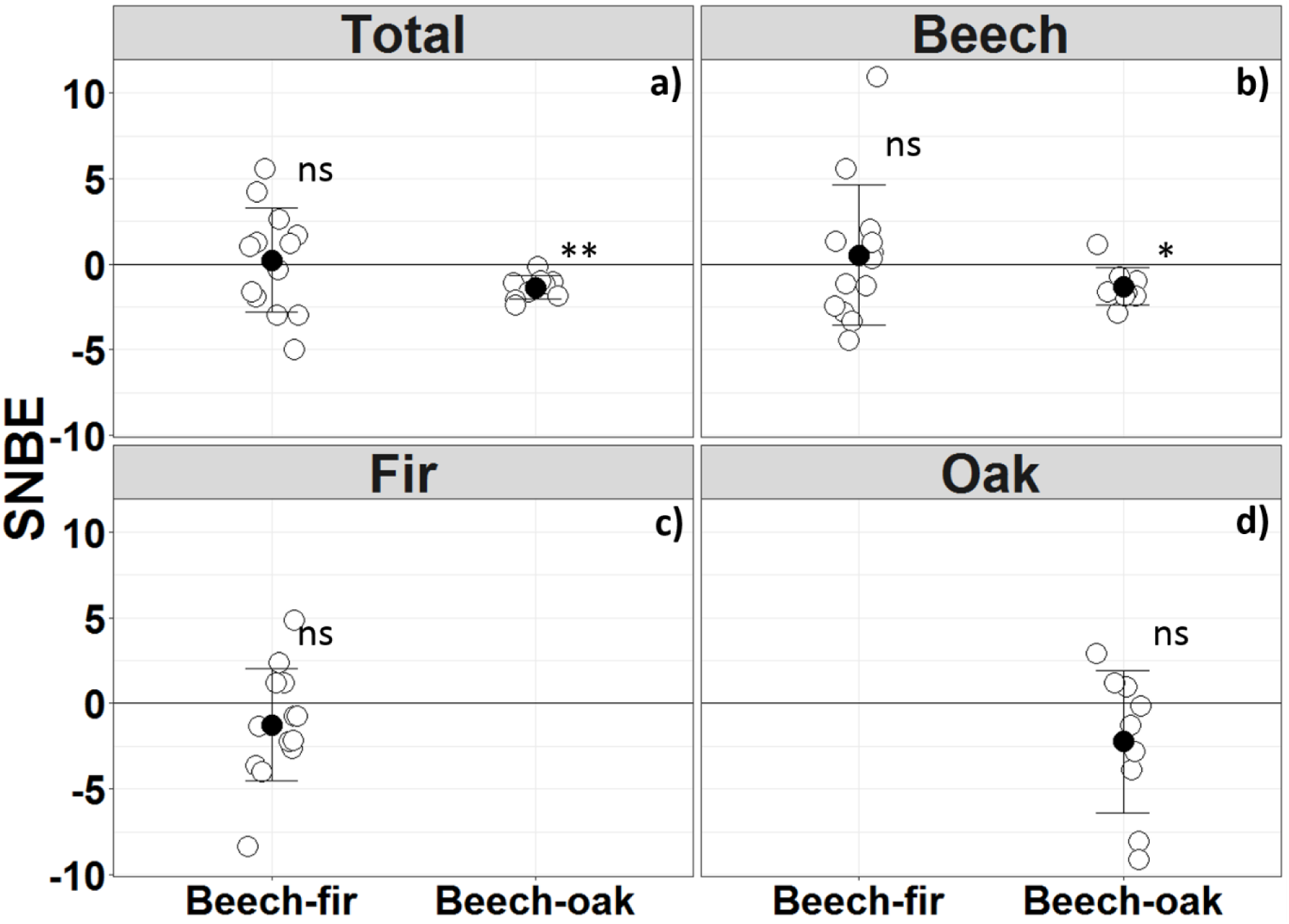
SNBE index values across the gradient. Each white point represents to a mixed stand and black dots represent the mean values (with deviation bars). A positive SNBE corresponds to a higher observed temporal stability than theoretical and a negative SNBE means SNBE corresponds a lower observed temporal stability than theoretical. The calculations of SNBE were done by considering all trees in the plots (“total”) [(a) n=19], only beech trees (b), only fir trees (c), and only oak trees (d). Beech-fir forest includes plots in Mont Ventoux, Vercors and Bauges and Beech-oak forests includes plots in Luberon Lagarde, Grand Luberon and Sainte-Baume. **: T-test’s p-value < 0.005; *: p-value < 0.05; ns: non-significant. (R version 3.4.4, package ggplot2)

**Figure 3.**
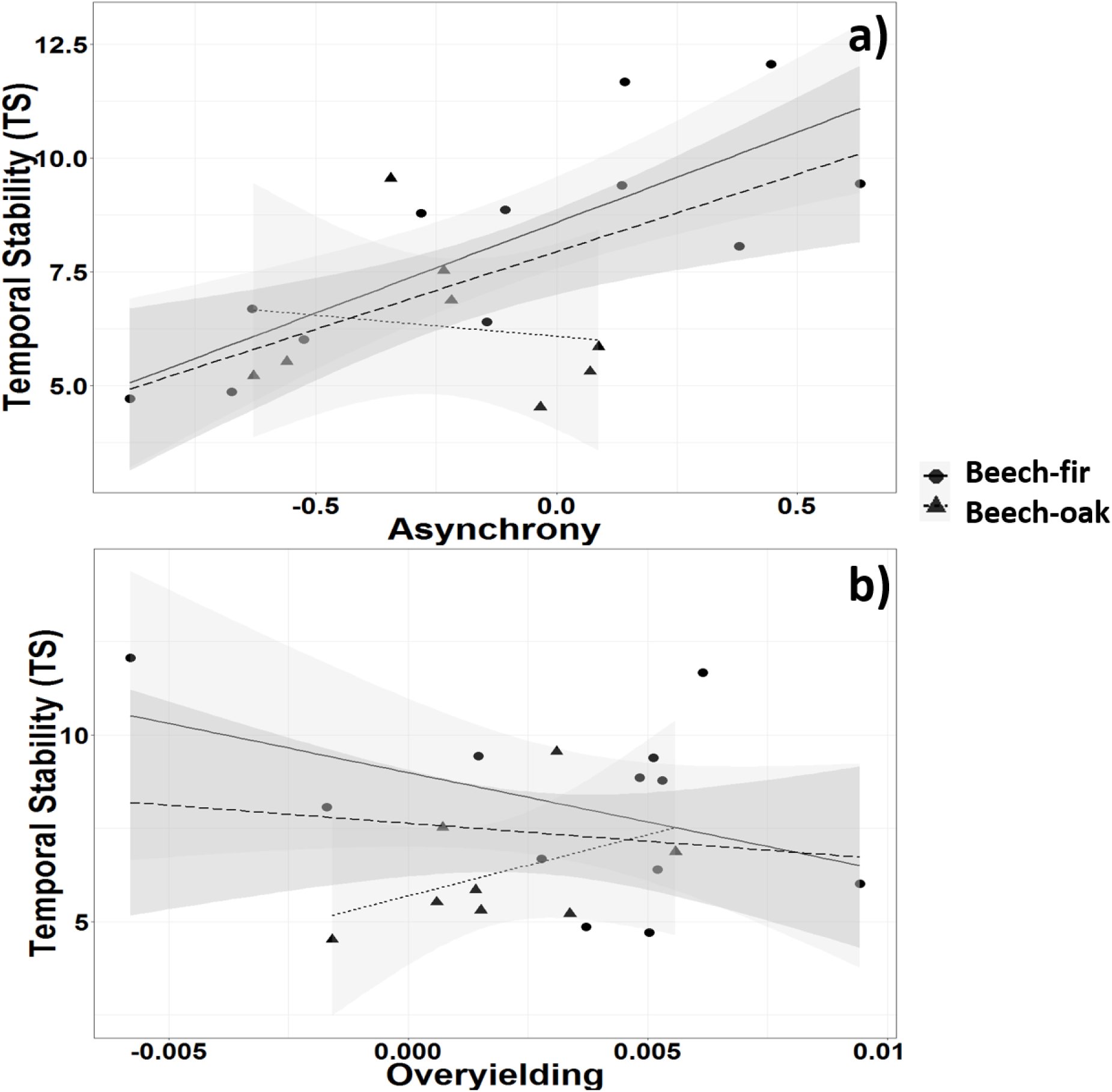
*TS* against asynchrony between species response (ɳ) and overyielding (*NBE*) across the gradient. a) *TS* against asynchrony. b) *TS* against *NBE*. The models either included only plots in the beech-fir forests (full line and circle), only the beech-oak forests (dotted line and triangle) or all plots across the gradient (long-dashed line). One symbol (circle or triangle) corresponds to one triplet; the lines are the regression line of each model. Models estimates can be found in Table 1. (R version 3.4.4, package ggplot2)

Considering the analysis of the effect of environmental conditions on TS, no trend has been depicted in the beech-fir forests (slope estimate = −0.07; P > 0.05, r² = 0.001, n= 13). Contrariwise the ratio between temporal stability of mixed and monospecific forests significantly decreased with increased productivity in the beech-oak forests (slope estimate = −1.16; P < 0.01, r² = 0.49, n=8), meaning that diversity effect on temporal stability of productivity decreased in favorable conditions (i.e. ratio inferior to 1, Fig. 5).

**Figure 4.**
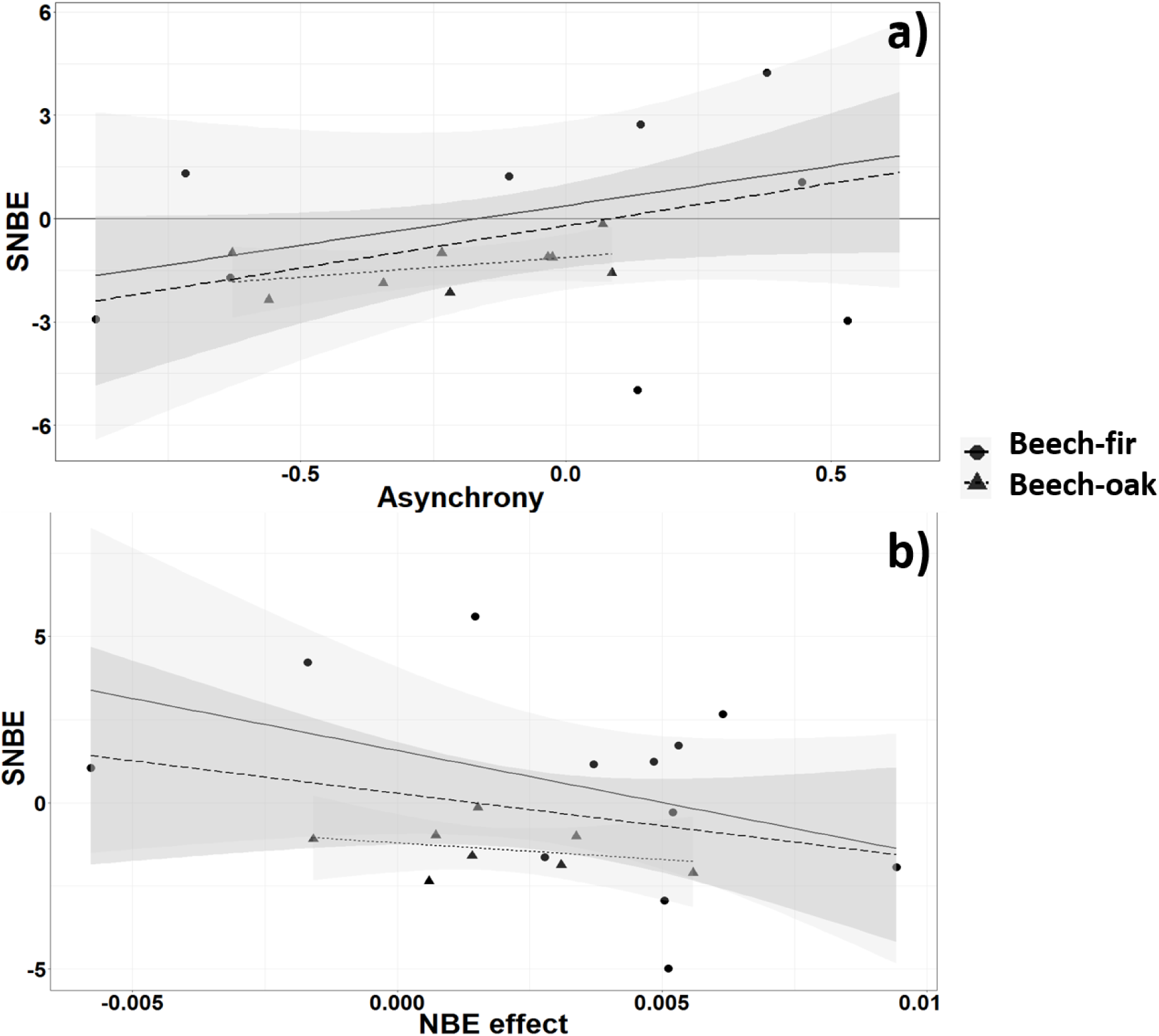
*SNBE* against asynchrony between species response (ɳ) and overyielding (*NBE*) across the gradient. a) *SNBE* against asynchrony. b) *SNBE* against *NBE*. The models either included only plots in the beech-fir forests (full line and circle), only the beech-oak forests (dotted line and triangle) or all plots across the gradient (long-dashed line). One symbol (circle or triangle) corresponds to one triplet; the lines are the regression line of each model. Models estimates can be found in Table 1. (R version 3.4.4, package ggplot2)

**Figure 5.**
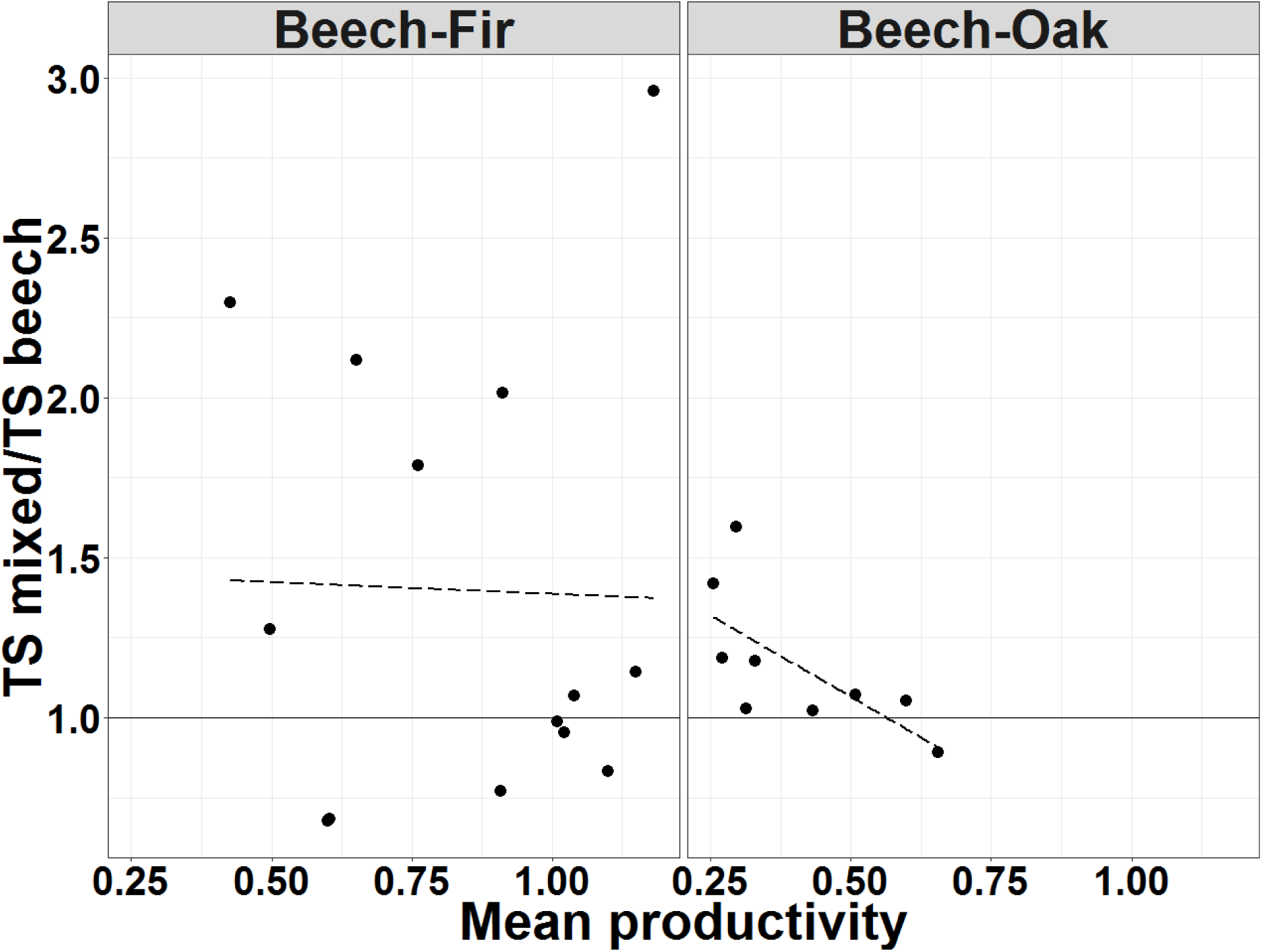
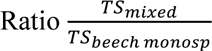 against mean productivity, using mean productivity as an integrative proxy for the sites’ conditions across the two gradients. Beech-fir forests include plots in Mont Ventoux, Vercors and Bauges (at left), and beech-oak forests include plots in Luberon Lagarde, Grand Luberon and Sainte-Baume (at right). (R version 3.4.4, package ggplot2)

### 3.2 SNBE

The *SNBE* index was either positive or negative in the beech-fir sites depending on the triplet. Contrariwise *SNBE* of mixed beech-oak plots was systematically negative (Fig. 2-a).

Calculating *SNBE* for each species separately, we found that a negative non-additive negative effect of diversity on productivity of beech trees in Southern sites (Fig. 2-b), while no trend was detected for oaks in Southern sites and for beech and fir in Northern sites (Fig. 2-b-c-d).

Focusing on mixed plots, we tested the effect of several potential drivers on *SNBE*. Relying on AIC to compare the several linear models, we found that *SNBE* was significantly dependent on asynchrony and *NBE* for the whole gradient (Table 2-a). In beech-fir and beech-oak forests, NBE and asynchrony have no significant effect on non-additive effect of diversity on temporal stability. According to the AIC criterion, *SNBE ∼ 1* (null model) is equivalent to *SNBE* ∼ ɳ.

No trend has been depicted regardless the dataset considered (P> 0.05) for the relationship between productivity and *SNBE* ratio.

We found the same trend with the first method (i.e. a significant positive effect of asynchrony on *SNBE*, but no significant effect of overyielding on *SNBE*) and the second method (i.e. with considering possible differences in total basal area between plots inside a triplet in *SNBE* calculation) regarding the effect of asynchrony on *SNBE* across the whole gradient, without any significant effect of climate on each kind of tested forests. It is noteworthy that the effect of overyielding on *SNBE* was on average negative, although not significant (Appendix 5 and 6).

## 4 DISCUSSION

We found that species richness only marginally (i.e. with p-value between 0.05 and 0.1) affected the *TS* in annual productivity for both kinds of mixed stands studied here. However, *TS* appeared to depend on asynchrony in species responses to environmental conditions (Table 1), especially for the beech-fir stands. The results on *SNBE*, depicting non-additive effects of species richness in the stability of productivity, showed that stability of mixed stands did not strongly differ from monospecific stands, as expected (Table 2). The variability in *SNBE* values is shown to be dependent on species composition, as *SNBE* was negative for oak-beech stands and not significant for fir-beech stands. There was also no difference in *SNBE* between trees in mixed or monospecific plots when each species was examined separately, except for beech in the beech-oak forests (Fig. 2).

The few studies testing the effect of stand diversity on temporal stability of productivity in forest ecosystems generally reported contrasted results (DeClerck et al. 2006; Jucker et al. 2014; del Río et al. 2017), as some of them showed increasing stability with stand diversity (Jucker et al. 2014; del Río et al. 2017) while others do not find any effect (DeClerck et al. 2006). Our results did not show strong differences between monospecific and mixed plots regarding TS, while they showed a decrease in *SNBE* for one of the mixtures tested and no significant effect for the other one. However, any comparisons with our results remain nevertheless limited, because the cited studies focused on different mixed stands than those studied here: conifer stands in Sierra Nevada (DeClerck et al. 2006) or Scots pine-beech across Europe (del Río et al. 2017). Jucker et al. (2014) tested diversity effect on *TS* in various forest types in six sites across Europe, including beech-fir stands in one site (Romania) but no specific results on these stands were provided, thus hampering a possible comparison with our own findings. However, although it is not enough to draw any generalities, it is noticeable that the negative effect of diversity on *TS* reported by Jucker et al. (2014) occurred in the only Mediterranean site tested, thus in similar conditions in which the negative effect was found in our study (beech-oak mixture).

Regarding the differences between the effect of diversity on stability in productivity at the community *vs.* species levels, (Tilman et al. 2006) showed that diversity may stabilize the productivity of the community while destabilizing the productivity of each species by analyzing experimental data on grasslands, reconciling an old debate about contrasted diversity effects at the community and species levels (Ives and Carpenter 2007). Although it focused on one specific two-species mixture (beech-Scot pine stands), (del Río et al. 2017) found consistent results with Tilman et al. (2006). Our results also confirmed this trend for beech-oak stands, especially at the species level, as mean *SNBE* values are significantly negative at species level for beech.

Our results further showed that asynchrony in species’ responses to environmental fluctuations increased *TS* of mixed beech-fir stands, while *NBE* and asynchrony did not affect significantly *SNBE* (although the effect of asynchrony was almost marginally significant) (Table 2). It is noticeable that our findings on the role of asynchrony on *TS* were consistent with theoretical expectations (Cardinale et al. 2013, but see Morin et al. 2014), and with other empirical studies on forest ecosystems (Jucker et al. 2014; del Río et al. 2017). Our results on *SNBE* tend to suggest that the trend highlighted for *TS* mostly relies on pure additive effects, i.e. related to dominance effects.

In several studies, it has been demonstrated that asynchrony in species responses leads to compensatory dynamics between species (Loreau 2010; Morin et al. 2014), supporting the biodiversity insurance hypothesis (Yachi and Loreau 1999). This finding is also consistent with the finding that diversity effect on *TS* varies among species and stand types (de Mazancourt et al. 2013). In fact, in the case of strong asynchrony, co-existing species do not experience an increase or decrease in productivity during the same years, which may explain why stand productivity is generally more stable in mixed stands relative to monospecific stands. Species asynchrony is thus expected to be stronger in communities composed of functionally different species, as these species are expected to show a weaker covariation in their response to climate fluctuations in non-limiting conditions, i.e. not stressful ones (Hector et al. 2010). For instance, mixed stands stability including beech and Scot pine trees - thus two species with large functional differences (deciduous *vs.* evergreen, shade-tolerant *vs.* pioneer species, drought sensitive *vs* resistant) – is found to increase with increasing asynchrony of these two species’ responses (del Río et al. 2017), although most of the studied mixed stands actually show synchronous *BAI* time-series. In our study, the relatively weak effect of asynchrony on TS and SNBE may be related to the fact that physiological difference between species do not induce a strong enough asynchrony. For example, even if it is a deciduous-evergreen mixed stand, the two species have similar drought sensitivity and shade-tolerance. This weak difference may not be enough to induce a systematically strong asynchrony level at the stand level.

Regarding beech-oak mixed stands, the asynchrony effect on stability in productivity is consistent. In fact, almost all mixed beech-oak stands show negative *SNBE* (−1.37±0.69) and asynchrony (−0.21±0.26) values, which thus confirms that mixed stands with more synchronous species responses leads to stands less stable than expected in terms of productivity (Fig. 3-a). This may first seem surprising because beech and pubescent oak have different ecological strategies, with contrasting shade tolerances and, more importantly, various responses to environmental conditions. In fact, pubescent oak is supposed to be more resistant to drought than beech, while beech is better adapted to cold conditions (Rameau et al. 1999). The three Mediterranean sites (particularly Grand Luberon and Luberon-Lagarde) are characterized by intense drought events (with particularly very little summer precipitation), but they are also located at relatively high elevation (from 750 m to 1150 m *a.s.l.*), with low temperatures in winter and spring. Therefore, while beech trees may be more sensitive to drought, the growth of pubescent oak trees may be strongly affected by air and soil temperatures as these trees are close to their elevational limit in the region (Rameau et al. 1999). As a result, the growth rates of the two species in the Mediterranean sites tend to co-vary more strongly over time compared to the beech-fir stands, as we observe in the studied plots, leading, on average, to a reduced asynchrony and to a weak or even negative effect of tree diversity on stand stability.

Testing for forest composition effect on productivity stability necessarily implies studying same species under various climate conditions. In our study, climatic range in which the same mixed stands can co-occur (i.e. same co-existing species) and can be comparable (i.e. controlling for other environmental factors, such as aspect or soil conditions) may be very constrained, which thus strongly limits the range of climatic conditions that could be explored. We therefore had to split our analyses involving the test of the effect of climatic conditions on *TS* between the beech-fir and beech-oak stands, to avoid any confounding effect of co-variation in climate conditions within species composition (Table 1-2). In doing so, no effect of the variation in climate conditions, temperature and precipitation on *SNBE* is found neither in beech-oak stands (in the beech-oak forests) nor in beech-fir stand (in fir-beech forests). The number of triplets sampled appeared to still be limiting regardless of the large field sampling effort they necessitated. One original goal of the present study was indeed to test for the effect of environmental conditions on the link between tree diversity and ecosystem functioning, which has been rarely done previously (del Río et al. 2017).

In the last two decades, study of *NBE* (Loreau and Hector 2001) on mean productivity was classically done in *BEF*-studies focusing on grasslands (Huston 1997; Marquard et al. 2009) and on forests (Morin et al. 2011; Grossman et al. 2017). Our study is the first (with del Río et al. 2017, to our knowledge, to assess the relationship between diversity and temporal stability of productivity in forests while controlling for species’ relative abundances between mixed and monospecific stands. Then, similarly to the use of *NBE* instead of comparing the mean productivity of mixed and monospecific stands (as illustrated in Morin et al. 2011), we believe that using the *SNBE* allows to focus on non-additive effect of diversity on stability in productivity, which in turn allows to better depict the diversity effect in mixed stands, especially as the relative abundance of species in mixed forests necessarily varied among and within sites. This standardization between mixed stands also allows a better exploration of the underlying processes driving temporal stability in productivity (e.g. overyielding and asynchrony). It also allows removing a large part of external variability, such as environmental conditions and variability in stand characteristics (e.g. structure) between triplets. This triplet-based design is thus perfectly suited to this kind of analysis.

Regarding the potential links between the SGH and the diversity effects on stability in productivity, we find results consistent with the SGH when considering the *TS* ratio between mixed stands and beech monospecific stands for beech-oak stands, but not for beech-fir stands. A possible explanation is that environmental conditions in beech-fir stands may not be harsh enough to highlight any effect of the SGH, as already discussed. Contrarily, *TS* increased in mixed beech-oak stands relatively to monospecific beech stands with increasing environmental harshness in the Mediterranean sites where the conditions are much more stressful than in the beech-fir forests. However, this pattern disappears when considering *SNBE*. This may suggest that environmental harshness modulates the diversity effect on stability in productivity mostly through changes in species dominance (i.e. a non-additive effect).

The forest plots considered here were mostly high forests. However, one site included coppice stands (Grand Luberon – although it is noticeable that there was no management in this site for the last 40 years at least). We did not consider this difference in stand structure because Grand Luberon was the most stressful site, and it was thus impossible to disentangle the two effects. However, this point must be kept in mind, especially to interpret the results in the Grand Luberon site.

Finally, we believe that the approach developed here is key to properly understand how climate variability and diversity may affect temporal stability in productivity, especially for forest ecosystems for which BEF-experiments are difficult to carry-out and monitor in the long-term. However, using such a design imposes intense field work and monitoring, bringing some constraints as mentioned above, which led us to focus on a restricted number of mixtures. A continuation of this work would be to extend the range of mixtures sampled, by focusing on mixed stands including species predicted to be sensitive to climate change in this region, e.g. *Pinus sylvestris* and *Picea abies*, in addition to the three species studied here. Another important criterion in the selection of the mixed stand composition should be the persistence or transience of the mixture (Cordonnier et al. 2018). The study of permanent, or long-lasting, mixtures, as done here, provides more relevant information on the mechanisms driving diversity-productivity relationships than more transient mixtures (e.g. beech – Scots pine mixed stands del Río et al. 2017) (Cordonnier et al. 2018).

## DATA ACCESSIBILITY

Data will be deposited in the Dryad Digital Repository.

## AUTHORS’ CONTRIBUTIONS

XM conceived the original question and field setup of this study. MJ and XM designed the research and developed the methodology. MJ, JB and XM collected the data; CP simulated the climatic data; MJ processed and analyzed the data; MJ and XM led the writing of the manuscript. All authors contributed critically to the drafts and gave final approval for publication.

## ACKNOWLEDGMENTS

This study strongly benefitted from the help of E. Defossez, for his assistance in collecting the data. We thank S. Coq and several students for additional help in collecting the data. T. Cordonnier, P. Vallet and F. Lebourgeois provided helpful comments on earlier versions of the manuscript. We also thank L. Gillespie who corrected the English of the manuscript. We greatly thank the *Office National des Forêts* for allowing access to the sites, and especially J. Ladier, P. Dreyfus and C. Riond from the *Recherche-Développement-Innovation* department for their help in selecting the plots. MJ benefitted from an ADEME grant. This study was mostly funded by the project DISTIMACC (ECOFOR-2014-23, French Ministry of Ecology and Sustainable Development, French Ministry of Agriculture and Forest) and benefitted from the ANR project BioProFor (contract no. 11-PDOC-030-01) and from the project REFORM (# 2816ERA02S, PCIN2017026) from the framework of Sumforest ERA-NET.

