## Supplementary Materials for "Tree diversity and the temporal stability of mountain forest productivity: testing the effect of species composition, through asynchrony and overyielding"

### TITLE

#### AUTHORS DETAILS

[1] CEFE UMR 5175, CNRS – Université de Montpellier – Université Paul-Valéry Montpellier – EPHE, 1919, route de Mende, F-34293 Montpellier cedex 5, France

[2] Agence de l'environnement et de la Maîtrise de l'Energie, 20, avenue du Grésillé- BP 90406 49004 Angers Cedex 01, France

[3] AgroParisTech, UMR1092, Laboratoire d'Étude des Ressources Forêt-Bois (LERFoB), ENGREF

14 rue Girardet, CS14216, FR- 54042 Nancy Cedex, France

### Appendix 1: Description of the sites.

| *Site* | *Coordinates (°)* | *MAT (°C)* | *SAP (mm)* | $N_{plot}$ *mixed* | $N_{plot}$ *beech* | $N_{plot}$ *fir* | $N_{plot}$ *oak* |
|---|---|---|---|---|---|---|---|
| Bauges (Combe d'Ire) | 45.697930°N 6.214553°E | 6.3-7.6 | 1994-2079 | 4 | 5 | 3 | - |
| Vercors (Lente) | 44.928504°N 5.321516°E | 5.4-7.1 | 1387-1541 | 5 | 5 | 4 | - |
| Mont Ventoux (Beaumont du Ventoux) | 44.187901°N 5.253608°E | 5.6-7.3 | 1208-1224 | 5 | 5 | 5 | - |
| Luberon Largarde (Lagarde d'Apt) | 43.973001°N 5.479875°E | 9.2-9.6 | 1026-1027 | 2 | 2 | - | 2 |
| Grand Luberon (Saint-Martin du Castillon) | 43.819412°N 5.535047°E | 9.7-10.7 | 790-796 | 3 | 3 | - | 3 |
| Sainte Baume | 43.334609°N 5.766041°E | 9.9-10.2 | 938-941 | 4 | 3 | - | 4 |

*Coordinates*: latitude and longitude; *MAT*: minimal and maximal of mean annual temperature; *SAP*: minimal and maximal sum of annual precipitation; $N_{plot}$ : number of sampled plots according to the forest type (i.e. mixed stand – *mixed* – and monospecific stand - *fir*, *beech* or *oak*).

### Appendix 2:

| | Bauges | Vercors | Ventoux | Luberon Lagarde | Grand Luberon | Ste-Baume |
|---|---|---|---|---|---|---|
| Beech | 0.30/0.67 | 0.30/0.41 | 0.45/0.50 | 0.41/0.64 | 0.29/0.46 | 0.26/0.34 |
| Fir | 0.60/0.58 | 0.34/0.36 | 0.53/0.37 | - | - | - |
| Oak | - | - | - | 0.16/0.49 | 0.51/0.31 | 0.26/0.16 |

Adjusted r-squares of linear models testing the effect of *G* on *BAI* for each site, each species in each stand (in mixed stand/in monospecific stand). Model: $BAI_j \sim G_j$ , with $j$ representing the year.

### Appendix 3:

| Site | Triplet | Variable | Monospecific beech stand | Monospecific oak/fir stand | Mixed stand |
|---|---|---|---|---|---|
| Bauges | A | $H_{max}$ | 34 | 35 | 34 |
| | | D | 541 | 605 | 668 |
| | | $G_{tot}$ | 51 | 70 | 83 |
| | | %coppice | 0 | 0 | 0 |
| | | $Gini_{DBH}$ | 0.37 | 0.39 | 0.38 |
| | | TS | 8.21 | 15.11 | 9.39 |
| | B | $H_{max}$ | 34 | 43 | 34 |
| | | D | 541 | 796 | 764 |
| | | $G_{tot}$ | 51 | 93 | 69 |
| | | %coppice | 0 | 0 | 0 |
| | | $Gini_{DBH}$ | 0.37 | 0.44 | 0.34 |
| | | TS | 8.21 | 4.94 | 8.78 |
| | C | $H_{max}$ | 37 | 31 | 33 |
| | | D | 637 | 700 | 573 |
| | | $G_{tot}$ | 85 | 90 | 74 |
| | | %coppice | 0.55 | 0 | 0 |
| | | $Gini_{DBH}$ | 0.46 | 0.46 | 0.38 |
| | | TS | 3.94 | 10.58 | 11.67 |
| Vercors | D | $H_{max}$ | 28 | 27 | 30 |
| | | D | 318 | 573 | 573 |
| | | $G_{tot}$ | 53 | 58 | 61 |
| | | %coppice | 0 | 0 | 0 |
| | | $Gini_{DBH}$ | 0.11 | 0.26 | 0.17 |
| | | TS | 4.95 | 7.63 | 8.86 |
| | E | $H_{max}$ | 27 | 27 | 31 |
| | | D | 668 | 573 | 637 |
| | | $G_{tot}$ | 47 | 58 | 86 |
| | | %coppice | 0 | 0 | 0 |
| | | $Gini_{DBH}$ | 0.21 | 0.26 | 0.34 |
| | | TS | 6.31 | 7.63 | 6.01 |
| | F | $H_{max}$ | 17 | 22 | 19 |
| | | D | 1114 | 955 | 1273 |
| | | $G_{tot}$ | 28 | 52 | 36 |
| | | %coppice | 0 | 0 | 0 |
| | | $Gini_{DBH}$ | 0.2 | 0.25 | 0.32 |
| | | TS | 8.02 | 7.12 | 6.69 |
| | G | $H_{max}$ | 34 | 30 | 27 |
| | | D | 446 | 446 | 509 |
| | | $G_{tot}$ | 65 | 65 | 47 |
| | | %coppice | 0 | 0 | 0 |
| | | $Gini_{DBH}$ | 0.2 | 0.21 | 0.2 |
| | | TS | 7.74 | 6.36 | 5.24 |
| | H | $H_{max}$ | 28 | 26 | 26 |
| | | D | 541 | 477 | 764 |
| | | $G_{tot}$ | 68 | 51 | 82 |
| | | %coppice | 0 | 0 | 0 |
| | | $Gini_{DBH}$ | 0.16 | 0.28 | 0.34 |

| | | | | | |
|---|---|---|---|---|---|
| | | TS | 5.21 | 3.13 | 4.85 |
| Ventoux | I | $H_{max}$ | 15 | 24 | 27 |
| | | D | 2355 | 1464 | 1719 |
| | | $G_{tot}$ | 45 | 75 | 71 |
| | | %coppice | 0.78 | 0 | 0.35 |
| | | $Gini_{DBH}$ | 0.16 | 0.3 | 0.32 |
| | | TS | 3.20 | 6.94 | 6.4 |
| | J | $H_{max}$ | 15 | 16 | 18 |
| | | D | 2355 | 541 | 1560 |
| | | $G_{tot}$ | 39 | 53 | 57 |
| | | %coppice | 0.53 | 0 | 0.57 |
| | | $Gini_{DBH}$ | 0.18 | 0.16 | 0.35 |
| | | TS | 6.31 | | 8.06 |
| | K | $H_{max}$ | 20 | 17 | 23 |
| | | D | 955 | 2483 | 1623 |
| | | $G_{tot}$ | 60 | 68 | 57 |
| | | %coppice | 0.53 | 0 | 0.41 |
| | | $Gini_{DBH}$ | 0.18 | 0.25 | 0.29 |
| | | TS | 5.25 | 7.97 | 12.05 |
| | L | $H_{max}$ | 22 | 19 | 23 |
| | | D | 764 | 509 | 1560 |
| | | $G_{tot}$ | 57 | 49 | 64 |
| | | %coppice | 0 | 0 | 0 |
| | | $Gini_{DBH}$ | 0.29 | 0.27 | 0.37 |
| | | TS | 6.10 | 7.22 | 4.71 |
| Grand Luberon | M | $H_{max}$ | 17 | 10 | 12 |
| | | D | 2005 | 2005 | 2165 |
| | | $G_{tot}$ | 39 | 29 | 29 |
| | | %coppice | 0.59 | 0.65 | 0.61 |
| | | $Gini_{DBH}$ | 0.2 | 0.15 | 0.2 |
| | | TS | 4.56 | 7.42 | 6.48 |
| | N | $H_{max}$ | 14 | 12 | 12 |
| | | D | 2324 | 1719 | 2642 |
| | | $G_{tot}$ | 38 | 44 | 36 |
| | | %coppice | 0.79 | 0.52 | 0.64 |
| | | $Gini_{DBH}$ | 0.19 | 0.18 | 0.17 |
| | | TS | 4.96 | 13.11 | 5.85 |
| | O | $H_{max}$ | 18 | 10 | 12 |
| | | D | 1719 | 1655 | 2292 |
| | | $G_{tot}$ | 29 | 22 | 32 |
| | | %coppice | 0.59 | 0.54 | 0.82 |
| | | $Gini_{DBH}$ | 0.15 | 0.17 | 0.15 |
| | | TS | 7.32 | 17.19 | 7.52 |
| Luberon Lagarde | P | $H_{max}$ | 20 | 13 | 16 |
| | | D | 732 | 1241 | 1910 |
| | | $G_{tot}$ | 44 | 29 | 47 |
| | | %coppice | 0 | 0.51 | 0.72 |
| | | $Gini_{DBH}$ | 0.38 | 0.24 | 0.21 |
| | | TS | 9.35 | 9.62 | 9.55 |
| | Q | $H_{max}$ | 19 | 13 | 15 |
| | | D | 1655 | 1178 | 3406 |
| | | $G_{tot}$ | 42 | 21 | 49 |
| | | %coppice | 0.81 | 0.59 | 0.81 |

| | | | | | |
|---|---|---|---|---|---|
| | | $Gini_{DBH}$ | 0.3 | 0.16 | 0.19 |
| | | TS | 7.71 | 17.18 | 6.87 |
| Ste Baume | R | $H_{max}$ | 29 | 19 | 21 |
| | | D | 1178 | 1910 | 1496 |
| | | $G_{tot}$ | 70 | 80 | 70 |
| | | %coppice | 0 | 0 | 0 |
| | | $Gini_{DBH}$ | 0.39 | 0.32 | 0.3 |
| | | TS | 2.83 | 10.83 | 4.52 |
| | S | $H_{max}$ | 21 | 17 | 21 |
| | | D | 1719 | 1178 | 2228 |
| | | $G_{tot}$ | 43 | 34 | 52 |
| | | %coppice | 0.33 | 0 | 0.3 |
| | | $Gini_{DBH}$ | 0.25 | 0.2 | 0.22 |
| | | TS | 4.96 | 6.92 | 5.21 |
| | T | $H_{max}$ | 21 | 16 | 19 |
| | | D | 1719 | 955 | 1019 |
| | | $G_{tot}$ | 43 | 38 | 34 |
| | | %coppice | 0.33 | 0 | 0 |
| | | $Gini_{DBH}$ | 0.25 | 0.21 | 0.22 |
| | | TS | 4.96 | 4.85 | 5.3 |

Table describing the structure of the stands, with *code_site*, $H_{max}$ (m), *D* (tree/ha), $G_{tot}$ (m²/ha), *%coppice* and $Gini_{DBH}$ respectively corresponding to plots ID, tree maximum height in the plot, tree density in the plot, total basal area of the plot, coppice proportion (ratio of coppice tree number and total tree number in one plot) and Gini index coefficient calculated on DBH, representing structural heterogeneity in DBH.

### Appendix 4:

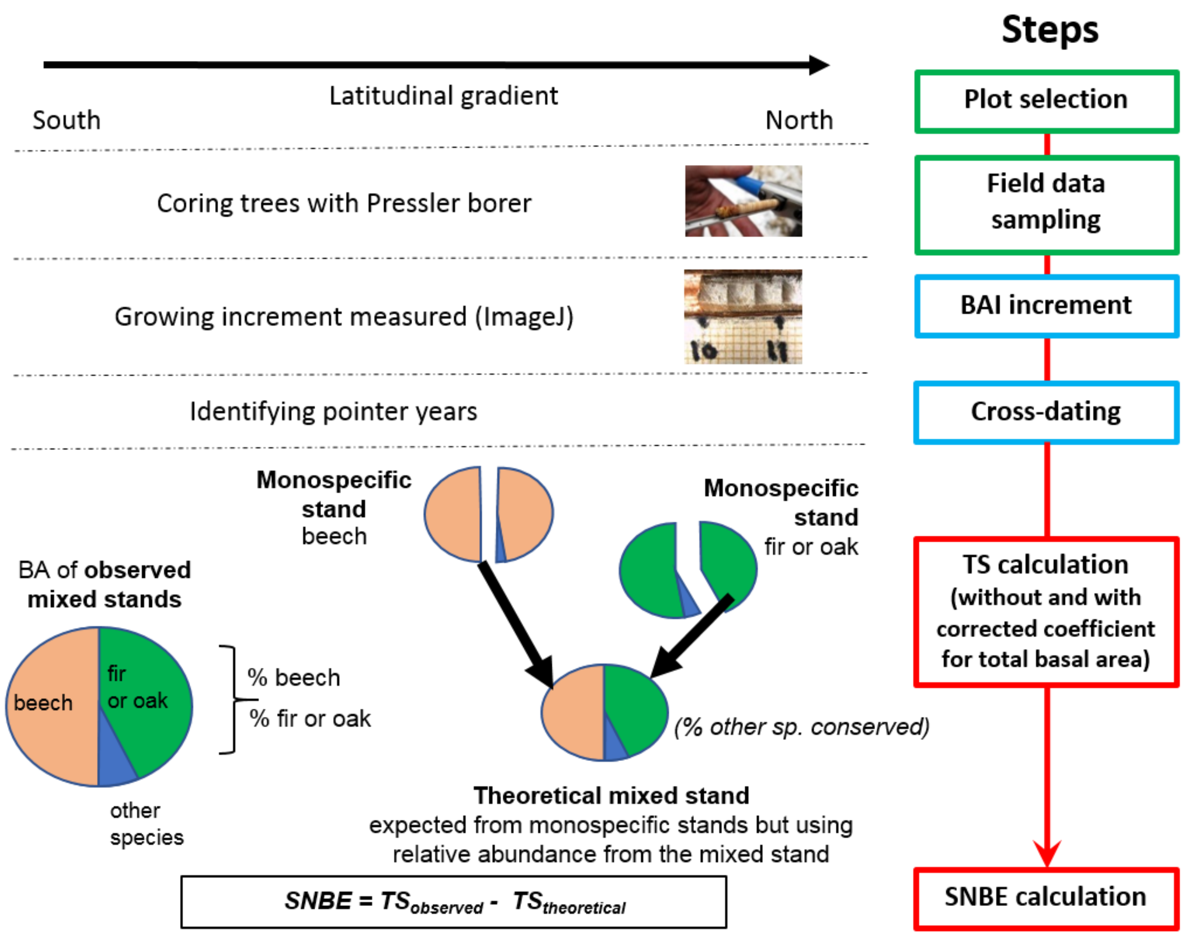


Description of the several steps of the dendrochronological analyses carried-out in the study to finally calculate the SNBE index.

### Appendix 5:

(a)

| Model | Total | beech-fir | beech-oak |
|---|---|---|---|
| *SNBE ~ 1* | 97 | 63 | **25** |
| *SNBE ~ η* | 89 | **60** | 24 |
| *SNBE ~ NBE* | 89 | 65 | 27 |
| *SNBE ~ η + NBE* | 81 | 62 | 25 |
| *SNBE ~ Composition* | 95 | - | - |
| *SNBE ~ η + Composition* | 87 | - | - |
| *SNBE ~ NBE + Composition* | 86 | - | - |
| *SNBE ~ Gtot + Composition* | 94 | - | - |
| *SNBE ~ η + NBE + Composition* | **79** | - | - |
| *SNBE ~ η + NBE + Gtot + Composition* | 78 | - | - |
| *SNBE ~* $T_{mean}$ *+ P* | - | 65 | 27 |
| *SNBE~ η + NBE +* $T_{mean}$ *+ P* | - | 63 | - |

(b)

| *Dataset* | *Variables* | *Estimate* | *SE* | $R^2$ |
|---|---|---|---|---|
| Total (n=22) | ***η*** | **3.31** | **1.16** | |
| | *NBE* | -6.22 | 73.63 | |
| | *Composition* | -1.52 | 0.94 | |
| beech-fir stands (n=13) | ***η*** | **3.58** | **1.53** | - |
| beech-oak stands (n=9) | - | - | - | - |

Linear mixed models of Net Biodiversity Effect on Stability (*SNBE*) against asynchrony (*η*), Net Biodiversity Effect (*NBE)* and climate variables for the whole dataset (*Total*), and composition are "beech-fir" in North and "beech-oak" in South. For beech-fir and beech-oak stands of the gradient taken separately, $T_{mean}$ and *P* are annual mean temperature average and annual precipitation average across 1997-2012 respectively. In every model *site* is a random effect. *η* represents number of triplets considered in each model.

(a) AIC the most relevant tested linear models. AIC of most parsimonious models are highlighted in bold. (b) Coefficients estimates of explanatory variables in the selected models. Significant explicative variables are in bold.

### Appendix 6:

(a)

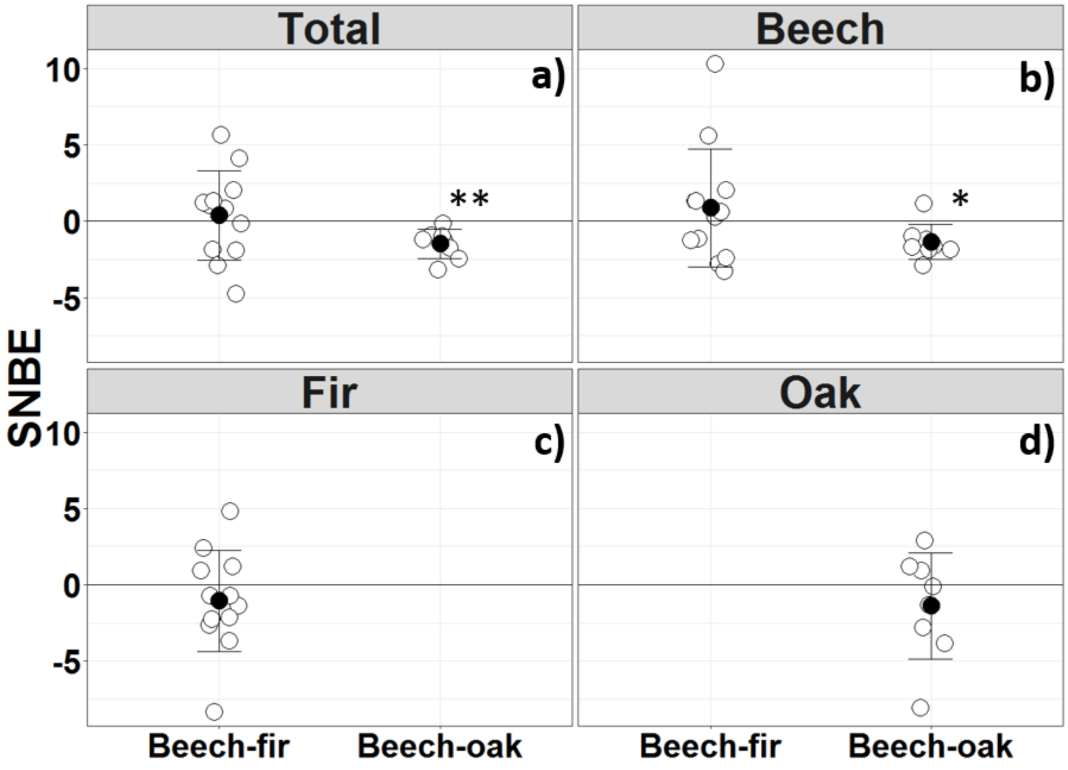


SNBE index values across the gradient. Open dots represent a mixed stand and black dots represent the mean values (with deviation bars). A positive SNBE corresponds to a stronger observed temporal stability than expected (under the null assumption of no effect of interspecific interactions on SNBE), and a negative SNBE corresponds to a weaker observed temporal stability than expected. The calculations of SNBE were done by considering all trees in the plots ("total") [(a) n=19], only beech trees (b), only fir trees (c), and only oak trees (d). Beech-fir forests include plots in Mont Ventoux, Vercors and Bauges and Beech-oak forests include plots in Luberon Lagarde, Grand Luberon and Sainte-Baume. **: T-test's p-value < 0.005; *: p-value < 0.05; ns: non-significant.

(b)

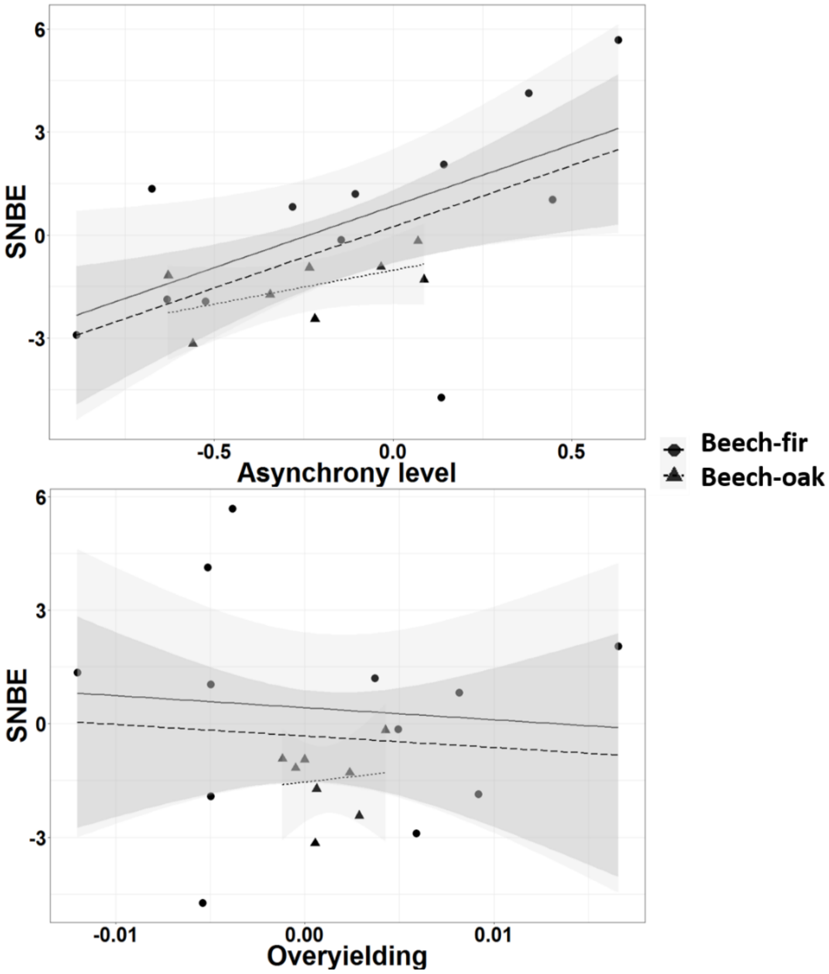


*SNBE* against asynchrony between species response ($\eta$) and overyielding (*NBE*) across the gradient. *SNBE* against asynchrony (at the top). *SNBE* against *NBE* (at the bottom). The models either included only plots in the beech-fir forests (full line and circle), only the beech-oak forests (dotted line and triangle) or all plots across the gradient (long-dashed line). One symbol (circle or triangle) corresponds to one triplet; the lines are the regression line of each model. Models estimates can be found in Table 1.
